# Asgard archaea are diverse, ubiquitous, and transcriptionally active microbes

**DOI:** 10.1101/374165

**Authors:** Mingwei Cai, Yang Liu, Zhichao Zhou, Yuchun Yang, Jie Pan, Ji-Dong Gu, Meng Li

**Affiliations:** Institute for Advanced Study, Shenzhen University, Shenzhen, China; Key Laboratory of Optoelectronic Devices and Systems of Ministry of Education and Guangdong Province, College of Optoelectronic Engineering, Shenzhen University, Shenzhen, China; Laboratory of Environmental Microbiology and Toxicology, School of Biological Sciences, The University of Hong Kong, Pokfulam Road, Hong Kong SAR, China

## Abstract

Asgard is a newly proposed archaeal superphylum. Phylogenetic position of Asgard archaea and its relationships to the origin of eukaryotes is attracting increasingly research interest. However, in-depth knowledge of their diversity, distribution, and activity of Asgard archaea remains limited. Here, we used phylogenetic analysis to cluster the publicly available Asgard archaeal 16S rRNA gene sequences into 13 subgroups, including five previously unknown subgroups. These lineages were widely distributed in anaerobic environments, with the majority of 16S rRNA gene sequences (92%) originating from sediment habitats. Co-occurrence analysis revealed potential relationships between Asgard, Bathyarchaeota, and Marine Benthic Group D archaea. Genomic analysis suggested that Asgard archaea are potentially mixotrophic microbes with divergent metabolic capabilities. Importantly, metatranscriptomics confirmed the versatile lifestyles of Lokiarchaeota and Thorarchaeota, which can fix CO_2_ using the tetrahydromethanopterin Wood-Ljungdahl pathway, perform acetogenesis, and degrade organic matters. Overall, this study broadens the understandings of Asgard archaea ecology, and also provides the first evidence to support a transcriptionally active mixotrophic lifestyle of Asgard archaea, shedding light on the potential roles of these microorganisms in the global biogeochemical cycling.

---

The three-domains-of-life tree theory divides cellular life into three domains: Bacteria, Archaea, and Eukarya^1^. Ever since Archaea became widely recognized as a separate domain of life, the exploration of the origin of Eukarya has intensified^2, 3^. Recently, metagenomic analyses revealed that Lokiarchaeota are the closest relative of Eukarya^4^. Its sister lineages, Thorarchaeota, Odinarchaeota, and Heimdallarchaeota, were then identified through metagenomic reconstruction^5-7^. Together, these lineages form the Asgard superphylum, elucidating the origin of eukaryotic cellular complexity^7^. However, some scientists argued the obtained Asgard genomes might be contaminated and the concatenated gene set for phylogenetic tree construction is not reliable^8, 9^. A recent study suggested that Asgard archaea might be non-phagocytotic, i.e., that the eukaryotic signature proteins (ESPs) of Asgard archaea did not originate in extracellular environments^10^, rendering their relationship with eukaryotes a mystery. Asgard archaea were initially identified in marine habitats, including lineages formerly named Marine Benthic Group B (MBG-B)^11^, Deep-Sea Archaeal Group (DSAG)^12^, Ancient Archaeal Group (AAG)^13^, and Marine Hydrothermal Vent Group (MHVG)^13, 14^. Identification of additional sequences revealed that these archaea inhabit various environments (e.g., lake sediment, mangrove sediment, estuarine sediment, and mud volcano)^5, 7, 15^. Indeed, Asgard archaea might be more diverse than the above-mentioned four phyla, as suggested by the recent discovered expressed Asgard-associated archaeal rRNA^15^. Thus, many questions remain to be answered regarding the diversity and classification of Asgard archaea, as well as the ecological distributions and the habitats of its lineages. In addition, current knowledge of the metabolic functions of Asgard archaea is mostly derived from the indirect inferences from metagenomic analysis. For example, genome reconstruction suggests that Lokiarchaeota might be anaerobic hydrogen-dependent autotrophs^16^, while Thorarchaeota might be mixotrophic^5^, and perhaps capable of acetogenesis and sulfur reduction^6^. However, the lack of cultured Asgard archaea representatives or culture-independent evidences (e.g., metatranscriptomics and DNA-SIP analysis) hampers a further understanding of the physiological functions and ecological roles of these archaea^6, 7^.

In the current study, we first used the publicly available 16S rRNA gene sequences to investigate the diversity and global distribution of Asgard archaea, as well as the relationship of Asgard subgroups with environmental factors and interactions with other archaeal lineages. Most importantly, we reconstructed and compared the potential metabolic pathways of Asgard archaea based on the publicly available genomes, and further confirmed their metabolic activities by metatranscriptomic analysis. To the best of our knowledge, this is the first-ever metatranscriptomic analysis of Asgard archaea that provides insights into their roles in the global biogeochemical cycling.

## Results and discussion

### Asgard archaea are highly diverse

Archaeal 16S rRNA gene sequences from two public databases (SILVA SSU 132 and NCBI databases) and sequences from a reference paper^15^ were utilized for a systematic investigation of Asgard archaea diversity. After sequence filtering, 451 operational taxonomic units (OTUs, 95% cutoff, 10,440 sequences) belonging to the Asgard archaeal superphylum were obtained. The majority of OTUs (79.4%) and sequences (85.1%) fell into the Lokiarchaeota lineage (Fig. 1 and Supplementary Table 1). According to the criteria of minimum intragroup similarity (>75%^17^) and minimum sequence number (>20) within a subgroup (see Methods), these sequences were divided into 13 subgroups (97.9% of all sequences) with high bootstrap values (>95%; Supplementary Table 1), including the known lineages Lokiarchaeota, Thorarchaeota, Odinarchaeota, and Heimdallarchaeota^7^. This indicates that the Asgard superphylum is more diverse than previously proposed^7^. For comparative purposes, DNA sequences retrieved from the databases (excluding expressed 16S rRNA gene sequences from the reference paper^15^) were used to reconstruct a phylogenetic tree (Supplementary Fig. 1). The tree topology was similar to that presented in Fig. 1 but with lower bootstrap values. Based on the tree phylogeny, five new Asgard subgroups (Asgard-1 to Asgard-5) were obtained, incorporating most sequences (99.7%) from the expressed 16S rRNA genes reported in the reference paper^15^. For analysis purposes, the broad Lokiarchaeota subgroup (minimum intragroup similarity of 80%; Supplementary Table 1) was subdivided into four adjacent groups (Loki-1, Loki-2a, Loki-2b, and Loki-3) based on the tree nodes (Fig. 1). MBG-B, previously considered to be Thorarchaeota^18^, were clustered with Lokiarchaeota (Supplementary Fig. 2). As the minimum intragroup similarity (71%, Supplementary Table 1) of sequences within Heimdallarchaeota was below the suggested phylum-level threshold of 75%^17^, the lineage was divided into two subgroups [Heimdall (AAG) and Heimdall (MHVG)], as previously recommended^7, 19^. Three of the 13 subgroups (Asgard 2–4) comprised sequences from the reference study^15^ (Supplementary Table 1), while only two lineages [Heimdall (AAG) and Loki-3] contained >70% sequences from public databases, indicating that the previously used universal archaeal primers or Asgard-specific primers^20-22^ retrieved sequences only from certain Asgard subgroups. We suggest that alternative approaches, e.g., RNA-extraction-based method^15^ or deep-sequencing technology^7^, will provide additional sequences for primer-set design, thus informing comprehensive and precise phylogenetic reconstruction of the Asgard superphylum in the future.

**Figure 1.**
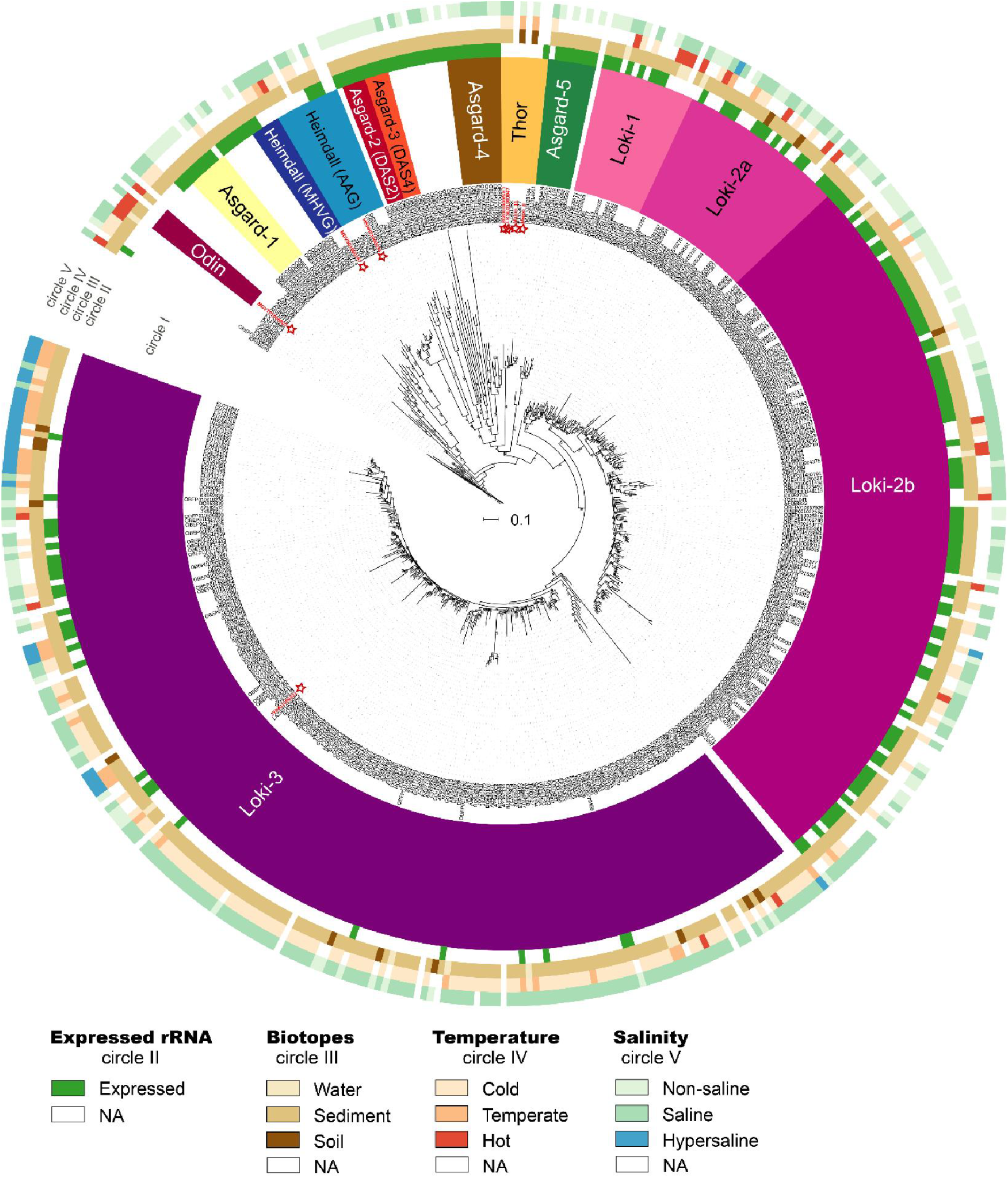
Maximum likelihood phylogenetic tree of Asgard archaea constructed using genome-based (red star) and publicly available 16S rRNA gene sequences clustered at 95% sequence identity. The tree was constructed with IQ-TREE using the maximum likelihood method based on 1000 bootstrap iterations and displayed using iTOL. Bootstrap support values >70 are shown. Subgroups were designated based on sequences fulfilling the sub-dividing criteria (see Methods). The outer color circles, from the inside to the outside, represent the expressed 16S rRNA, habitat, temperature, and salinity, accordingly (see Supplementary Table 3 for more details).

### Asgard archaea are ubiquitous and co-occur with other archaeal lineages

Current knowledge of the Asgard archaea habitats mainly comes from the extant Asgard genomes extracted from anaerobic sedimentary environments (Supplementary Table 2). In the current study, global investigations of the sequences of Asgard 16S rRNA genes deposited in public datasets indicated that a high percentage (~92%) of OTUs (95% cutoff) originated from sediment samples (Figs. 1 and 2a, Supplementary Table 3), similarly to Bathyarchaeota^23, 24^. In addition, Asgard archaea are thought to inhabit other potentially anaerobic environments, such as the marine water column, mud volcano, and soil (Figs. 1 and 2a). Detection of the genes (e.g., *mvh*, *frh*, and *nfn*^16, 25^, Supplementary Table 4) involving in anaerobic metabolisms corroborated the notion that Asgard archaea are anaerobic microbes. The smallest subcluster Odin, and portions of the Loki-1 and Loki-2a subgroups contained sequences retrieved from hydrothermal habitats; OTUs collected from hypersaline habitats (e.g., hypersaline microbial mat and salt-works belt) were clustered together in the Loki-3 subgroup (Fig. 1), suggesting similar subgroup biotopes. Since the investigated libraries were almost all recovered from offshore or marine sediments (Fig. 2a and Supplementary Table 3), most OTUs (~65%) have been found in cold saline environments (Fig. 1).

**Figure 2.**
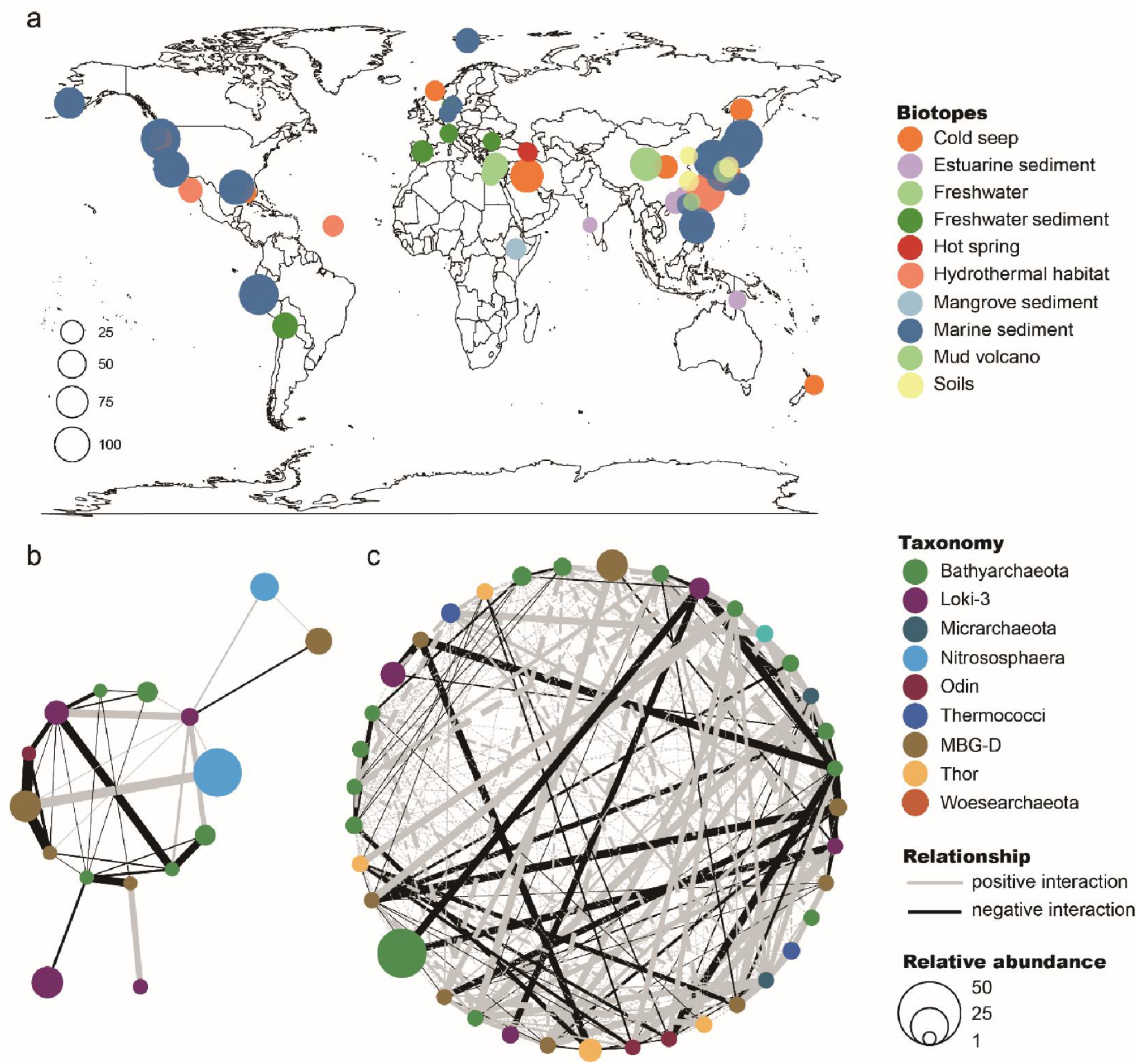
Global distribution of Asgard archaea and their co-occurrence with other archaeal lineages. **a** Global distribution and biotopes of Asgard archaea from 172 public archaeal libraries. Detailed information can be found in Supplementary Table 3. **b** Co-occurrence network of archaeal lineages from 144 public libraries. Nodes represent 90% OTUs and node size corresponds to the relative abundance of the Asgard 16S rRNA gene sequences. **c** Co-occurrence network of archaeal lineages from 110 mangrove and intertidal mudflat sediment samples. Nodes represent 97% OTUs and node size corresponds to the relative abundance of the Asgard 16S rRNA gene sequences. In **b** and **c**, lines represent interactions, with the line thickness corresponding to the corrected coefficient values calculated by Cytoscape (default correlation coefficient threshold and *p* < 0.05).

To investigate the relative abundance of Asgard sequences in different biotopes, we selected 65 published archaeal 16S rRNA libraries (Supplementary Table 3) for analysis. The analysis indicated that Asgard sequences in the selected samples accounted for up to 100% of the archaeal 16S rRNA libraries (Supplementary Fig. 3), with a higher percentage of Asgard sequences in deep marine sediments, especially the methane hydrate subseafloor sediments^14, 25^, than in soils or freshwater environments. Four of the 13 subgroups [i.e., Heimdall (AAG), Loki-2a, Loki-2b, and Loki-3 subgroups] were represented in these samples, with Loki-3 being the predominate subgroup in most cases (Supplementary Fig. 3). This extremely high abundance of Asgard archaea, especially the Loki-3 subgroup, indicated that they might be the keystone species for biogeochemical cycling in the anaerobic, cold, salty, and high-pressure environments.

16S rRNA gene sequences from public libraries (172 samples) and mangrove sediment samples (143 samples) obtained in the current study were utilized to investigate the relationships between archaea. Co-occurrence analysis indicated that the Asgard lineages Loki-3, Thor, and Odin shared the highest non-random association with Bathyarchaeota and Marine Benthic Group D (MBG-D) (Figs. 2b and 2c). Potential correlations within Asgard OTUs were also observed (e.g., Odin with Thor). Notably, Asgard OTUs from the same subgroup (e.g., Loki-3) exhibited different co-occurrence patterns with certain non-Asgard archaeal OTUs (Figs. 2b and 2c), suggesting potential divergent metabolic capabilities within the same Asgard subgroup. Further, Asgard archaea might co-occur with bacteria. However, the lack of bacterial data hampered in-depth investigations. Systematic investigations of the relative abundance and co-occurrence of archaea and bacteria, using appropriate primer sets^26^ (e.g., 515F–806R) are highly recommended for future studies.

### Metabolic differences among the Asgard phyla

To verify the metabolic divergence revealed by the co-occurrence analysis, publicly available genomes (Supplementary Fig. 4 and Supplementary Table 2) were used for metabolic capability comparisons, including the unreported pathways of Odinarchaeota and Heimdallarchaeota^27^. Genomic data suggested that all Asgard phyla might lead mixotrophic lifestyles utilizing both organic and inorganic carbon sources (e.g., protein, glucose, ethanol, and CO_2_). These analyses also indicated possible sulfur and nitrogen metabolisms (Fig. 3 and Supplementary Fig. 5). However, discrepancies between the phyla were more apparent than similarities, as revealed by KEGG module comparison (Supplementary Fig. 6), especially for Odinarchaeota and/or Heimdallarchaeota (Fig. 3). In the Wood-Ljungdahl pathway (WL; the reductive acetyl-CoA pathway), Lokiarchaeota and Thorarchaeota may use both tetrahydrofolate (THF) and tetrahydromethanopterin (THMPT) as C_1_ carriers, as has been proposed for methanogenic archaea^28^. By contrast, Heimdallarchaeota and Odinarchaeota appeared to reduce inorganic carbon by using one of the C_1_ carriers through THF-WL or THMPT-WL pathway, respectively. Typically, THF is a C_1_ carrier in bacterial acetogens^29^ and some halophilic archaea^30^. Although Heimdallarchaeota have been mainly identified in saline environments (Fig. 1), their ability to reduce CO_2_ via the THF-WL pathway required verification. Only Heimdallarchaeota harbored the complete gene set for the forward and reverse tricarboxylic acid (TCA) cycle (Fig. 3). Therefore, TCA cycle may not be the major energy-generating pathway for other Asgard lineages, similar to Hadesarchaea^31^. Furthermore, unlike the other three phyla, Odinarchaeota genomes are relatively small (1.5 Mbp, Supplementary Table 2), and lack many genes for butyryl-CoA oxidation and sulfate reduction (Fig. 3). Odinarchaeota, Heimdallarchaeota, and Lokiarchaeota lack most of the key genes of the Calvin-Benson-Bassham cycle (CBB; the reductive pentose phosphate cycle) (Fig. 3). Within the CBB pathway, the conversion of ribulose-5P to ribulose-1,5P2 is an ATP-consuming process, while the WL pathway can generate energy^32^. Thus, it is likely that the WL pathway is the preferred carbon fixation pathway within the Asgard superphylum, especially in deep marine sediments where the energy supply is extremely low^33^. Similarly to other archaeal lineages of Hadesarchaea^31^ and Novel Archaeal Group 1^34^, genes encoding ATP-dependent hexokinase (*hk*) and glucokinase (*pfk*) responsible for the initial step of the glycolytic pathway were not detected in Odinarchaeota and Thorarchaeota. This suggested that these microbes might not utilize extracellular glucose and other sugars, despite the fact that sugar transporters are encoded in their genomes (Supplementary Table 5). Because of the limited genome information for Lokiarchaeota, Odinarchaeota, and Heimdallarchaeota, their potential metabolic capabilities require further investigations.

**Figure 3.**
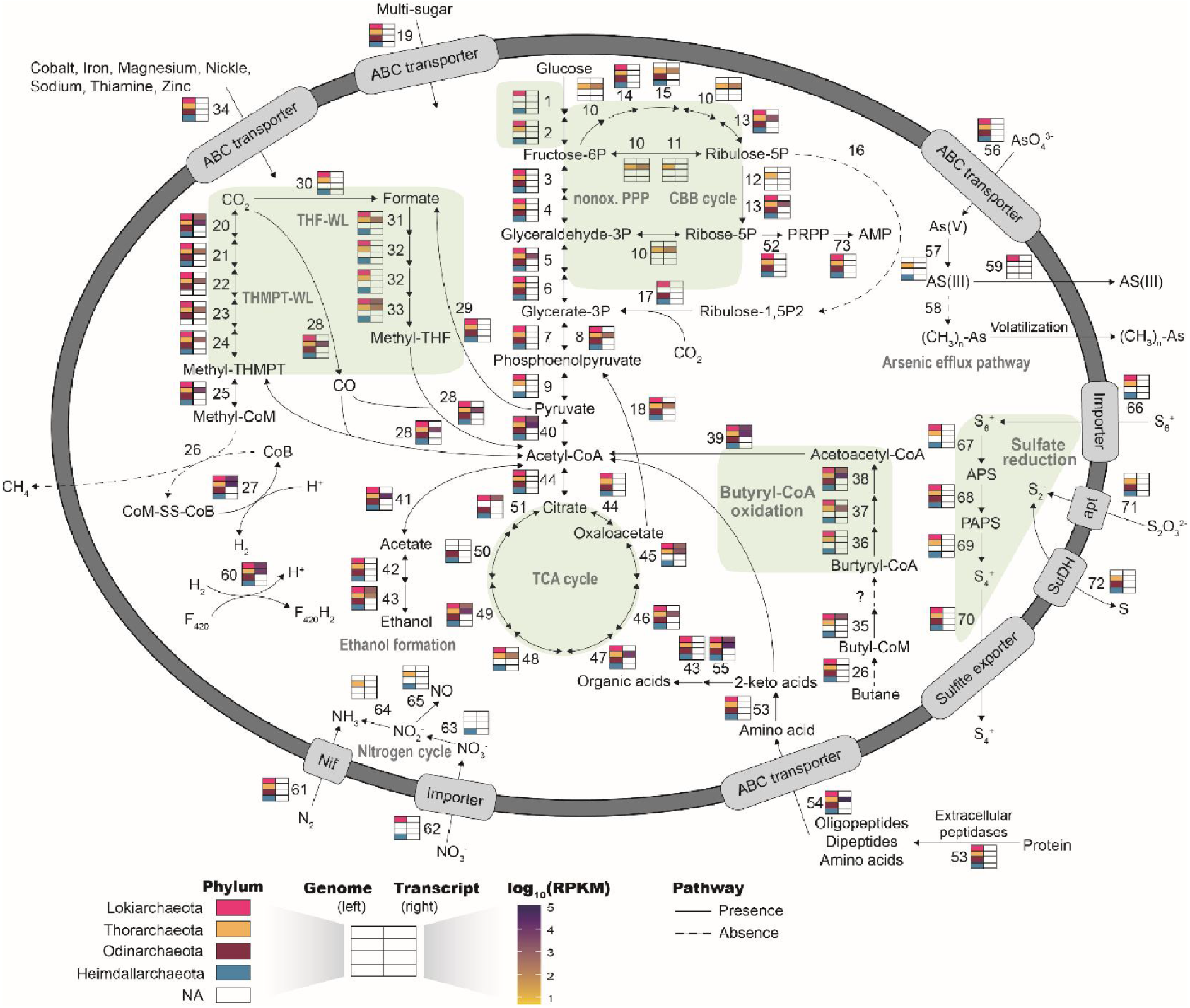
Metabolic differences among the Asgard phyla. The pathway was constructed using the publicly available Asgard genomes (Supplementary Table 2). Left portions of the rectangles represent genes from the Asgard genomes; right portions represent transcript abundances. Rectangles with different colors represent different phyla. Phyla exhibting metabolic differences were highlighted by green background. Detailed metabolic information for the genomes and transcripts is available in Supplementary Fig. 5. Pathway abbreviations: nonox. PPP, non-oxidative pentose phosphate pathway; CBB cycle, Calvin-Benson-Bassham cycle; THF-WL, tetrahydrofolate Wood-Ljungdahl pathway; THMPT-WL, tetrahydromethanopterin Wood-Ljungdahl pathway; TCA cycle, tricarboxylic acid cycle.

### Asgard archaea are transcriptionally active in the natural environment

To more precisely delineate the above-described metabolic capabilities of Asgard archaea, we combined metatranscriptomics with genome data for transcriptomic activity analysis (Fig. 3). Since most of the 16S rRNA gene sequences of Asgard archaea have been retrieved from offshore and deep marine environments (Fig. 1), the publicly available metatranscriptome datasets from these habitats were downloaded for BLASTn search, in addition to the metatranscriptome datasets for mangrove and intertidal mudflat sediments generated in the course of the current study (Supplementary Table 6). The analysis indicated that Lokiarchaeota and Thorarchaeota were transcriptionally active in offshore, mangrove, or intertidal mudflat sediments. By contrast, no transcripts originating from Heimdallarchaeota or Odinarchaeota were detected, leaving their metabolic activities unknown. This may be because of the high diversity of Asgard archaea (Fig. 1) and the currently limited availability of Asgard genome sequences (Supplementary Table 2). A complete transcript set for the THMPT-WL pathway confirmed the previous deduction for Thorarchaeota^5^, but the key gene (*fdh*) for the initial step of the THF-WL pathway (CO_2_ conversion to formate) was not expressed (Fig. 3). Transcripts involved in proteins and peptides degradation were present and abundant, indicating the proteins and/or peptides are major carbon and energy sources for Lokiarchaeota and Thorarchaeota. The highly expressed *acs* gene suggested that acetate could be one of the major organic carbon sources of Thorarchaeota. Since acetate plays an important role in subsurface sulfate reduction and methanogenesis^35^, the reversible reaction (acetate conversion to acetyl-CoA) might be a major pathway supplying acetate to other anaerobic microbes, as required. Transcripts for genes involved in ethanol formation and butyryl-CoA oxidation were also identified in Lokiarchaeota and Thorarchaeota. Intriguingly, no transcripts for nitrogen fixation, sulfate reduction, and arsenic efflux were detected. Further exploration of the Asgard transcriptomes may help to understand their metabolic activities in the sulfur and nitrogen cycle. We also detected expressed ESP genes of the Lokiarchaeota and Thorarchaeota lineages, especially the highly expressed genes for the conserved lokiactins and eukaryotic RLC7 family protein (Supplementary Fig. 7 and Supplementary Table 7), supporting the hypothesis that eukaryotes might have emerged from the Asgard superphylum^7^.

In conclusion, in the current study, we provided the first framework for the diversity, topology, global distribution, and activity of Asgard archaea, and their co-occurrence with other archaeal lineages. Biotope and genomic-based analyses indicated that these lineages are strictly anaerobic and engage in a mixotrophic lifestyle, although with different metabolic capacities. Moreover, metatranscriptomic evidence indicated that Lokiarchaeota and Thorarchaeota use the THMPT-WL pathway to fix CO_2_, perform acetogenesis, and/or degrade organic matter. To better understand the diversity and metabolism of Asgard archaea, further analyses, including metagenomics, metatranscriptomics, and experimental verification, are needed. This will undoubtedly provides fundamentally insights into the ecological roles of these microbes in the global biogeochemical cycling.

## Methods

### Data set construction

Archaeal 16S rRNA gene sequences were retrieved from the GenBank NCBI nucleotide database (September 2017) and SILVA SSU r132 database^36^. E-utilities^37^ was applied to search and retrieve the archaeal 16S rRNA gene sequences from the NCBI nucleotide database using the ESearch function with the following string: “16S AND 800:2000[Sequence Length] AND archaea[organism] AND rrna[Feature Key] AND isolation_source[All fields] NOT genome NOT chromosome NOT plasmid”. EFetch function was then used to retrieve the sequences and the corresponding GenBank-formatted flat file, which contained environmental information (e.g., location and isolation source). In the subsequent steps, custom scripts were designed to combine the two datasets and to remove low-quality (i.e., containing ‘N’ or shorter than 800 bp) and duplicate sequences, resulting in 100,786 archaeal sequences. To retain Asgard-associated sequences, these sequences were BLAST-searched against genome-based 16S rRNA sequences of Asgard archaea (≥800bp, Supplementary Table 1) using BLASTn with a cutoff E-value ≤1e-5, sequence identity ≥90%, and coverage ≥50%. This resulted in a set of 9765 potential Asgard sequences from the public databases. OTUs were assigned using the QIIME UCLUST 38 wrapper, with a threshold of 95% nucleotide sequence identity, and the cluster centroid for each OTU was chosen as the representative OTU sequence.

In addition to public databases, archaeal 16S rRNA gene sequences were also obtained from a recent study^15^. All expressed 16S rRNA gene sequences in the reference paper were BLAST-searched against a custom database containing 16S rRNA gene sequences retrieved from Asgard draft genomes and potential Asgard OTUs obtained as described above, with a cutoff E-value ≤1e-5, sequence identity ≥75%, and coverage ≥80%. Thus, 5588 potential Asgard sequences were obtained, including eight newly proposed clusters DAS1–8 (79 sequences)^15^. OTU re-formation of these archaeal sequences (15,353 sequences, from databases and the reference paper) was performed using the QIIME UCLUST^38^ wrapper, with a threshold of 95% pair-wise nucleotide sequence identity (1836 OTUs).

### Phylogenetic analysis of Asgard archaea

SINA-aligned^39^ representative OTU sequences obtained after dataset construction were pre-filtered using a backbone tree, which was constructed based on updated 16S rRNA gene datasets^27, 40^, using the ARB software (version 5.5) with the “Parsimony (Quick Add Marked)” tool^41^. This resulted in 462 Asgard-associated OTUs. Candidate representative OUT sequences (462 OTUs) and genome-based 16S rRNA gene sequences ≥600 bp (470 sequences in total) were used for phylogenetic tree construction. Maximum-likelihood tree was inferred with IQ-TREE (version 1.6.1)^42^ using the GTR+I+G4 mixture model (recommended by the “TESTONLY” model) and ultrafast (-bb 1000). Asgard subgroup designations were made when either a subgroup was defined in previous publications, or when subgroups with >20 sequences and similarity >75%^17^ were monophyletic. Attributes (i.e., expressed rRNA gene, biotope, temperature, and salinity) for each representative OTU were characterized and visualized using iTOL software^43^.

### Distribution of Asgard archaea

The corresponding environmental information (i.e., location and biotopes) for the 16S rRNA sequences of Asgard archaea was extracted from the GenBank-formatted flat file using custom scripts. This resulted in 172 libraries, 65 of which have been published (Supplementary Table 3). The relative abundance of Asgard archaea reported in the 65 studies was obtained by summarizing the archaeal histograms, archaeal tables, and/or sequence numbers in the archaeal phylogenetic trees. Information on the global distributing of sample locations, relative abundance, and subgroup diversity obtained for analyses was plotted using the mapdata, ggplot2, and scatterpie packages in R software.

### Sample collection and processing

Mangrove and intertidal mudflat sediment (totally 143 samples) were collected from the Shenzhen Bay^44^ (in press). DNA extraction, archaeal 16S rRNA gene sequence amplification, and high-throughput sequencing were performed as previously reported^44^ (in press). The 16S rRNA gene datasets were analyzed using QIIME 1 software^45^ and the sequences were clustered at 97% similarity threshold, resulting in 77,585 OTUs. OTUs with >1% relative abundance were chosen for downstream analysis.

Five samples (Maipo-7, Maipo-8, Maipo-9, Maipo-10, and Maipo-11) collected at different locations (mangrove and intertidal mudflat sediments) and different layer depths were obtained from the Mai Po Nature Reserve, Hong Kong, in 2017^5^. Details of the physicochemical parameters are specified elsewhere^5^. Shortly after sample collection, RNA was extracted, purified, and quantified, as described elsewhere^46^. The rRNA genes were removed from the total RNA using the Ribo-Zero rRNA removal kit (Illumina, Inc., San Diego, CA, USA) and reverse-transcribed. The cDNA was sequenced using an Illumina HiSeq sequencer (Illumina) with 150-bp paired-end reads, generating a total of 52.3-Gbp clean sequence data without adapters. Unassembled metatranscriptomic reads were quality-trimmed using Sickle^47^ with the quality score ≥25, and the potential rRNA reads were removed using SortMeRNA (version 2.0)^48^ against both SILVA 132 database and default databases (E-value cutoff ≤1e-5).

### Co-occurrence network construction

Relationships between Asgard OTUs and other archaeal OTUs were inferred based on an undirected co-occurrence network analysis using the CoNet tool in Cytoscape software (version 3.6.1)^49^. 16S rRNA archaeal OTUs from the public libraries (90% similarity cutoff) and mangrove sediments (97% similarity cutoff) were used in relationship analyses. Libraries with more than five OTUs were selected for network construction. The correlation coefficient thresholds were set using combined methods (e.g., Spearman correlation, Pearson correlation and Bray-Curtis Distance), as recommended by the manual^50^, and the Benjamini-Hochberg adjusted *p*-value < 0.05. The OTU taxonomy was annotated by searching against a custom database incorporating SILVA SSU 132 rRNA database and the Asgard sequences confirmed in the current study.

### Reconstruction of metabolic pathways

Publicly available Asgard genome sequences (Supplementary Table 1) were retrieved from the NCBI database for metabolic pathway reconstructing. Previously published Asgard genomes with low completeness^51^ were not included in the analysis. Protein-coding regions were predicted using Prodigal (version 2.6.3) and the “-p meta” option^52^. The KEGG server (BlastKOALA)^53^, eggNOG-mapper^54^, InterProScan tool (V60)^55^, and BLASTp vs. NCBI-nr database searched on December 2017 (E-value cutoff ≤1e-5) were used to annotate the protein-coding regions. Metabolic pathways were reconstructed based on the predicted annotations and the reference pathways depicted in KEGG and MetaCyc^56^. ESP prediction was based on arCOG (eggNOG-mapper) and the eukaryote-specific IPR domains (InterProScan)^7^ in Asgard genomes. Archaeal extracellular peptidases were identified using PRED-SIGNAL^57^ and PSORTb^58^ (Supplementary Table 8).

Metatranscriptome data from mangrove sediments and mudflat sediments, and other publicly available metatranscriptome data (Supplementary Table 6) were analyzed to clarify the transcriptomic activity of Asgard archaea. The abundance of transcripts for each gene from the Asgard archaeal genomes was determined by mapping all non-rRNA transcripts to predicted genes using BLASTn (E-value cutoff ≤1e-10). The BLAST hit with the highest score for each read was used for expression abundance analysis^59^. If mapped reads covered >90 genes in a genome, the genome was included in transcriptomic analyses. Normalized expression was calculated using the RPKM method (RPKM=[number of reads]/[transcript length, in kb]/[million mapped reads])^60^.

## Data availability

All archaeal 16S rRNA sequences were retrieved from NCBI database, SILVA SSU r132 database, and a reference paper as described in the Methods section. All Asgard genome sequences were from NCBI database. The metatranscriptomic data (Maipo-7, Maipo-8, Maipo-9, Maipo-10, and Maipo-11) are available in NCBI database with the accession number SRP150114.

## Author contributions

MC and ML conceived this study and wrote the manuscript. MC analyzed the 16S rRNA data and metatranscriptomic data. YL and ZZ provided suggestions for data analyses. ZZ and YY provided the mangrove metatranscriptomic data. ZZ and JP provided the mangrove 16S rRNA gene sequencing data. J-DG reviewed and improved the quality of manuscript.

## Acknowledgements

We greatly acknowledge all the authors who provided valuable data for this study. This research was funded by the National Natural Science Foundation of China (No. 31622002, 31600093, 31700430□ and 41506163), the Science and Technology Innovation Committee of Shenzhen (Grant No. JCYJ20170818091727570), and the Key Project of Department of Education of Guangdong Province (No.2017KZDXM071).

## References

1. Woese, C. R. and Fox, G. E., Phylogenetic structure of the prokaryotic domain: the primary kingdoms. Proc. Natl. Acad. Sci. USA 74, 5088–5090 (1977).

2. Pühler, G., Leffers, H., Gropp, F., Palm, P., Klenk, H.-P., Lottspeich, F., Garrett, R. A. and Zillig, W., Archaebacterial DNA-dependent RNA polymerases testify to the evolution of the eukaryotic nuclear genome. Proc. Natl. Acad. Sci. USA 86, 4569–4573 (1989).

3. Embley, T. M. and Martin, W., Eukaryotic evolution, changes and challenges. Nature 440, 623 (2006).

4. Spang, A., Saw, J. H., Jørgensen, S. L., Zaremba-Niedzwiedzka, K., Martijn, J., Lind, A. E., van Eijk, R., Schleper, C., Guy, L. and Ettema, T. J., Complex archaea that bridge the gap between prokaryotes and eukaryotes. Nature 521, 173–179 (2015).

5. Liu, Y., Zhou, Z., Pan, J., Baker, B. J., Gu, J.-D. and Li, M., Comparative genomic inference suggests mixotrophic lifestyle for Thorarchaeota. ISME J. 12, 1021 (2018).

6. Seitz, K. W., Lazar, C. S., Hinrichs, K. U., Teske, A. P. and Baker, B. J., Genomic reconstruction of a novel, deeply branched sediment archaeal phylum with pathways for acetogenesis and sulfur reduction. ISME J. 10, 1696–1705 (2016).

7. Zaremba-Niedzwiedzka, K., Caceres, E. F., Saw, J. H., Backstrom, D., Juzokaite, L., Vancaester, E., Seitz, K. W., Anantharaman, K., Starnawski, P., Kjeldsen, K. U., Stott, M. B., Nunoura, T., Banfield, J. F., Schramm, A., Baker, B. J., Spang, A. and Ettema, T. J., Asgard archaea illuminate the origin of eukaryotic cellular complexity. Nature 541, 353–358 (2017).

8. Da Cunha, V., Gaia, M., Gadelle, D., Nasir, A. and Forterre, P., Lokiarchaea are close relatives of Euryarchaeota, not bridging the gap between prokaryotes and eukaryotes. PLoS Genet. 13, e1006810 (2017).

9. Da Cunha, V., Gaia, M., Nasir, A. and Forterre, P., Asgard archaea do not close the debate about the universal tree of life topology. PLoS Genet. 14, e1007215 (2018).

10. Burns, J. A., Pittis, A. A. and Kim, E., Gene-based predictive models of trophic modes suggest Asgard archaea are not phagocytotic. Nat. Ecol. Evol. 2, 697 (2018).

11. Vetriani, C., Jannasch, H. W., MacGregor, B. J., Stahl, D. A. and Reysenbach, A.-L., Population structure and phylogenetic characterization of marine benthic archaea in deep-sea sediments. Appl. Environ. Microbiol. 65, 4375–4384 (1999).

12. Inagaki, F., Takai, K., Komatsu, T., Kanamatsu, T., Fujioka, K. and Horikoshi, K., Archaeology of Archaea: geomicrobiological record of Pleistocene thermal events concealed in a deep-sea subseafloor environment. Extremophiles 5, 385–392 (2001).

13. Takai, K. and Horikoshi, K., Genetic diversity of archaea in deep-sea hydrothermal vent environments. Genetics 152, 1285–1297 (1999).

14. Inagaki, F., Suzuki, M., Takai, K., Oida, H., Sakamoto, T., Aoki, K., Nealson, K. H. and Horikoshi, K., Microbial communities associated with geological horizons in coastal subseafloor sediments from the Sea of Okhotsk. Appl. Environ. Microbiol. 69, 7224–7235 (2003).

15. Karst, S. M., Dueholm, M. S., McIlroy, S. J., Kirkegaard, R. H., Nielsen, P. H. and Albertsen, M., Retrieval of a million high-quality, full-length microbial 16S and 18S rRNA gene sequences without primer bias. Nat. Biotechnol. 36, 190–195 (2018).

16. Sousa, F. L., Neukirchen, S., Allen, J. F., Lane, N. and Martin, W. F., Lokiarchaeon is hydrogen dependent. Nat. Microbiol. 1, 16034 (2016).

17. Yarza, P., Yilmaz, P., Pruesse, E., Glöckner, F. O., Ludwig, W., Schleifer, K.-H., Whitman, W. B., Euzéby, J., Amann, R. and Rosselló-Móra, R., Uniting the classification of cultured and uncultured bacteria and archaea using 16S rRNA gene sequences. Nat. Rev. Microbiol. 12, 635 (2014).

18. Adam, P. S., Borrel, G., Brochier-Armanet, C. and Gribaldo, S., The growing tree of Archaea: new perspectives on their diversity, evolution and ecology. ISME J. 11, 2407–2425 (2017).

19. Eme, L., Spang, A., Lombard, J., Stairs, C. W. and Ettema, T. J., Archaea and the origin of eukaryotes. Nat. Rev. Microbiol. 15, 711–723 (2017).

20. Fischer, M. A., Güllert, S., Neulinger, S. C., Streit, W. R. and Schmitz, R. A., Evaluation of 16S rRNA gene primer pairs for monitoring microbial community structures showed high reproducibility within and low comparability between datasets generated with multiple archaeal and bacterial primer Pairs. Front. Microbiol. 7, 1297 (2016).

21. Yu, Y., Lee, C., Kim, J. and Hwang, S., Group-specific primer and probe sets to detect methanogenic communities using quantitative real-time polymerase chain reaction. Biotechnol. Bioeng. 89, 670–679 (2005).

22. Ogawa, D. M., Moriya, S., Tsuboi, Y., Date, Y., Prieto-da-Silva, Á. R., Rádis-Baptista, G., Yamane, T. and Kikuchi, J., Biogeochemical Typing of Paddy Field by a Data-Driven Approach Revealing Sub-Systems within a Complex Environment-A Pipeline to Filtrate, Organize and Frame Massive Dataset from Multi-Omics Analyses. PLoS One 9, e110723 (2014).

23. Fillol, M., Auguet, J. C., Casamayor, E. O. and Borrego, C. M., Insights in the ecology and evolutionary history of the Miscellaneous Crenarchaeotic Group lineage. ISME J. 10, 665–77 (2016).

24. Zhou, Z., Pan, J., Wang, F., Gu, J.-D. and Li, M., Bathyarchaeota: globally distributed metabolic generalists in anoxic environments. FEMS Microbiol. Rev., (2018).

25. Inagaki, F., Nunoura, T., Nakagawa, S., Teske, A., Lever, M., Lauer, A., Suzuki, M., Takai, K., Delwiche, M. and Colwell, F. S., Biogeographical distribution and diversity of microbes in methane hydrate-bearing deep marine sediments on the Pacific Ocean Margin. Proc. Natl. Acid. Sci. USA 103, 2815–2820 (2006).

26. Peiffer, J. A., Spor, A., Koren, O., Jin, Z., Tringe, S. G., Dangl, J. L., Buckler, E. S. and Ley, R. E., Diversity and heritability of the maize rhizosphere microbiome under field conditions. Proc. Natl. Acad. Sci. USA 110, 6548–6553 (2013).

27. Spang, A., Caceres, E. F. and Ettema, T. J., Genomic exploration of the diversity, ecology, and evolution of the archaeal domain of life. Science 357, eaaf3883 (2017).

28. Buchenau, B. and Thauer, R. K., Tetrahydrofolate-specific enzymes in Methanosarcina barkeri and growth dependence of this methanogenic archaeon on folic acid or p-aminobenzoic acid. Arch. Microbiol. 182, 313–325 (2004).

29. Matthews, R. G. and Drummond, J. T., Providing one-carbon units for biological methylations: mechanistic studies on serine hydroxymethyltransferase, methylenetetrahydrofolate reductase, and methyltetrahydrofolate-homocysteine methyltransferase. Chem. Rev. 90, 1275–1290 (1990).

30. Boroujerdi, A. F. and Young, J. K., NMR-derived folate-bound structure of dihydrofolate reductase 1 from the halophile Haloferax volcanii. Biopolymers 91, 140–144 (2009).

31. Baker, B. J., Saw, J. H., Lind, A. E., Lazar, C. S., Hinrichs, K.-U., Teske, A. P. and Ettema, T. J., Genomic inference of the metabolism of cosmopolitan subsurface Archaea, Hadesarchaea. Nat. Microbiol. 1, 16002 (2016).

32. Borrel, G., Adam, P. S. and Gribaldo, S., Methanogenesis and the Wood-Ljungdahl pathway: an ancient, versatile, and fragile association. Genome Biol. Evol. 8, 1706–1711 (2016).

33. Magnabosco, C., Ryan, K., Lau, M. C., Kuloyo, O., Lollar, B. S., Kieft, T. L., Van Heerden, E. and Onstott, T. C., A metagenomic window into carbon metabolism at 3 km depth in Precambrian continental crust. ISME J. 10, 730 (2016).

34. Becraft, E. D., Dodsworth, J., Murugapiran, S. K., Thomas, S. M., Ohlsson, I., Stepanauskas, R., Hedlund, B. P. and Swingley, W. D., Genomic Comparison of Two Family-Level Groups of the Uncultivated NAG1 Archaeal Lineage from Chemically and Geographically Disparate Hot Springs. Front. Microbiol. 8, 2082 (2017).

35. Oremland, R. S. and Polcin, S., Methanogenesis and sulfate reduction: competitive and noncompetitive substrates in estuarine sediments. Appl. Environ. Microbiol. 44, 1270–1276 (1982).

36. Quast, C., Pruesse, E., Yilmaz, P., Gerken, J., Schweer, T., Yarza, P., Peplies, J. and Glöckner, F. O., The SILVA ribosomal RNA gene database project: improved data processing and web-based tools. Nucleic Acids Res. 41, D590-D596 (2012).

37. Kans, J., Entrez direct: E-utilities on the UNIX command line. (2017).

38. Edgar, R. C., Search and clustering orders of magnitude faster than BLAST. Bioinformatics 26, 2460–2461 (2010).

39. Pruesse, E., Peplies, J. and Glöckner, F. O., SINA: accurate high-throughput multiple sequence alignment of ribosomal RNA genes. Bioinformatics 28, 1823–1829 (2012).

40. Durbin, A. M. and Teske, A., Archaea in organic-lean and organic-rich marine subsurface sediments: an environmental gradient reflected in distinct phylogenetic lineages. Front. Microbiol. 3, 168 (2012).

41. Ludwig, W., Strunk, O., Westram, R., Richter, L., Meier, H., Yadhukumar Buchner, A., Lai, T., Steppi, S. and Jobb, G., ARB: a software environment for sequence data. Nucleic Acids Res. 32, 1363–1371 (2004).

42. Nguyen, L.-T., Schmidt, H. A., von Haeseler, A. and Minh, B. Q., IQ-TREE: a fast and effective stochastic algorithm for estimating maximum-likelihood phylogenies. Mol. Biol. Evol. 32, 268–274 (2014).

43. Letunic, I. and Bork, P., Interactive Tree Of Life (iTOL): an online tool for phylogenetic tree display and annotation. Bioinformatics 23, 127–128 (2006).

44. Zhou, Z., Meng, H., Liu, Y., Gu, J.-D. and Li, M., Stratified bacterial and archaeal community in mangrove and intertidal wetland mudflats revealed by high throughput 16S rRNA gene sequencing. Front. Microbiol. 8, (2017).

45. Caporaso, J. G., Kuczynski, J., Stombaugh, J., Bittinger, K., Bushman, F. D., Costello, E. K., Fierer, N., Pena, A. G., Goodrich, J. K. and Gordon, J. I., QIIME allows analysis of high-throughput community sequencing data. Nat. Methods 7, 335 (2010).

46. Li, M., Baker, B. J., Anantharaman, K., Jain, S., Breier, J. A. and Dick, G. J., Genomic and transcriptomic evidence for scavenging of diverse organic compounds by widespread deep-sea archaea. Nat. Commun. 6, 8933 (2015).

47. Joshi, N. and Fass, J., Sickle: A sliding-window, adaptive, quality-based trimming tool for FastQ files (Version 1.33)[Software]. (2011).

48. Kopylova, E., Noé, L. and Touzet, H., SortMeRNA: fast and accurate filtering of ribosomal RNAs in metatranscriptomic data. Bioinformatics 28, 3211–3217 (2012).

49. Shannon, P., Markiel, A., Ozier, O., Baliga, N. S., Wang, J. T., Ramage, D., Amin, N., Schwikowski, B. and Ideker, T., Cytoscape: a software environment for integrated models of biomolecular interaction networks. Genome Res. 13, 2498–2504 (2003).

50. Faust, K. and Raes, J., CoNet app: inference of biological association networks using Cytoscape. F1000Research 5, (2016).

51. Parks, D. H., Rinke, C., Chuvochina, M., Chaumeil, P.-A., Woodcroft, B. J., Evans, P. N., Hugenholtz, P. and Tyson, G. W., Recovery of nearly 8,000 metagenome-assembled genomes substantially expands the tree of life. Nat. Microbiol. 2, 1533 (2017).

52. Hyatt, D., Chen, G. L., LoCascio, P. F., Land, M. L., Larimer, F. W. and Hauser, L. J., Prodigal: prokaryotic gene recognition and translation initiation site identification. BMC Bioinformatics 11, 119 (2010).

53. Kanehisa, M., Sato, Y. and Morishima, K., BlastKOALA and GhostKOALA: KEGG tools for functional characterization of genome and metagenome sequences. J. Mol. Biol. 428, 726–731 (2016).

54. Huerta-Cepas, J., Forslund, K., Coelho, L. P., Szklarczyk, D., Jensen, L. J., von Mering, C. and Bork, P., Fast genome-wide functional annotation through orthology assignment by eggNOG-mapper. Mol. Biol. Evol. 34, 2115–2122 (2017).

55. Jones, P., Binns, D., Chang, H.-Y., Fraser, M., Li, W., McAnulla, C., McWilliam, H., Maslen, J., Mitchell, A. and Nuka, G., InterProScan 5: genome-scale protein function classification. Bioinformatics 30, 1236–1240 (2014).

56. Caspi, R., Foerster, H., Fulcher, C. A., Kaipa, P., Krummenacker, M., Latendresse, M., Paley, S., Rhee, S. Y., Shearer, A. G. and Tissier, C., The MetaCyc Database of metabolic pathways and enzymes and the BioCyc collection of Pathway/Genome Databases. Nucleic Acids Res. 36, D623-D631 (2007).

57. Bagos, P., Tsirigos, K., Plessas, S., Liakopoulos, T. and Hamodrakas, S., Prediction of signal peptides in archaea. Protein Eng. Des. Sel. 22, 27–35 (2008).

58. Yu, N. Y., Wagner, J. R., Laird, M. R., Melli, G., Rey, S., Lo, R., Dao, P., Sahinalp, S. C., Ester, M. and Foster, L. J., PSORTb 3.0: improved protein subcellular localization prediction with refined localization subcategories and predictive capabilities for all prokaryotes. Bioinformatics 26, 1608–1615 (2010).

59. Lesniewski, R. A., Jain, S., Anantharaman, K., Schloss, P. D. and Dick, G. J., The metatranscriptome of a deep-sea hydrothermal plume is dominated by water column methanotrophs and lithotrophs. ISME J. 6, 2257 (2012).

60. Robinson, M. D. and Oshlack, A., A scaling normalization method for differential expression analysis of RNA-seq data. Genome Biol. 11, R25 (2010).

